# Fibroblast Mechanotransduction Network Predicts Targets for Mechano-Adaptive Infarct Therapies

**DOI:** 10.1101/2020.08.13.250001

**Authors:** Jesse D. Rogers, William J. Richardson

**Affiliations:** Department of Bioengineering; Clemson University; Clemson, SC, 29631; USA

**Keywords:** cardiac fibroblast, mechanotransduction, fibrosis, signaling network, myocardial infarct, systems biology

## Abstract

Regional control of fibrosis after myocardial infarction is critical for maintaining structural integrity in the infarct while preventing collagen accumulation in non-infarcted areas. Cardiac fibroblasts modulate matrix turnover in response to biochemical and biomechanical cues, but the complex interactions between signaling pathways confounds efforts to develop therapies for regional scar formation. Here, we employed a logic-based ordinary differential equation model of fibroblast mechano-chemo signal transduction to predict matrix protein expression in response to canonical biochemical stimuli and mechanical tension. Functional analysis of mechano-chemo interactions showed extensive pathway crosstalk with tension amplifying, dampening, or reversing responses to biochemical stimuli. Comprehensive drug target screens in low- and high-tension contexts identified 13 mechano-adaptive therapies that promote matrix accumulation in regions where it is needed and reduce matrix levels in regions where it is not needed. Our predictions demonstrate this approach’s utility for discovering context-specific mechanisms mediating fibrosis and druggable targets for spatially resolved therapies.

## Introduction

Controlling cardiac fibrosis remains a major challenge in developing long-term treatments for patients suffering from myocardial infarction (MI). For a proportion of the over 800,000 patients diagnosed per year (Benjamin et al., 2019), excessive hypertrophy and scar formation in the non-infarcted myocardium can result in the development of heart failure, with decreases in contractility, myocardial conductivity, and pump function leading to poor rates of survival (Shah et al., 2017). Currently prescribed therapeutics for MI patients are designed to reduce hypertrophy via several neurohormonal mechanisms including inhibition of angiotensin II (AngII) and norepinephrine (NE) or increasing the bioavailability of natriuretic peptides (NPs) (Ponikowski et al., 2016), but no drugs have been approved for treating cardiac fibrosis directly. Notably, scar formation serves a reparative function in post-MI wound healing by replacing necrotic myocardium and preventing infarct expansion and cardiac rupture (Weber et al., 2013). Therefore, developing therapeutics that limit excessive fibrosis to prevent the development of heart failure while preserving beneficial scar tissue at the infarct site would be a critical step towards improving long-term patient outcomes.

Tissue remodeling in the myocardium is a dynamic process in which the accumulation of extracellular matrix proteins (e.g. collagens I and III, fibronectin) and matricellular proteins are balanced by degradation via proteases such as matrix metalloproteinases (MMPs). This balance is mediated largely by cardiac fibroblasts, which infiltrate the infarct site and assume an activated, synthetic phenotype to promote matrix accumulation (Chen and Frangogiannis, 2013). A wide variety of biochemical factors regulate fibroblast behavior such as AngII, transforming growth factor-β (TGFβ1), endothelin-1 (ET1), and various inflammatory cytokines (Leask, 2010; Turner, 2014), but a growing body of research has demonstrated that biomechanical factors such as circumferential tension and tissue stiffness are also critical determinants of fibroblast behavior (Saucerman et al., 2019). Parallel transduction of biochemical and biomechanical stimuli is integrated by fibroblasts through a complex signaling network in which pathways interact via systems-level crosstalk and feedback mechanisms (Hinz, 2015; Saraswati et al., 2020), making prediction and control of fibroblast behavior in the post-MI environment highly challenging.

Computational approaches for investigating cell signaling have shown promise for discovering mechanisms-of-action underlying pathological states and drug treatments. Previous models representing cardiac cell signaling as a network of interactions have proven successful in generating detailed insight into cardiomyocyte contraction and hypertrophy as well as fibroblast chemotransduction and various other biological systems (Saez-Rodriguez et al., 2015; Saucerman and McCulloch, 2004; Tan et al., 2017; Zeigler et al., 2020). Previous models have simulated cardiac and adventitial fibroblast activation and pro-fibrotic activity in response to biochemical and biomechanical cues, although limited exploration of mechanotransduction pathways in these models still leave unanswered questions about the role of tension in regional matrix turnover (Wang et al., 2020; Zeigler et al., 2016). In this current study, we expanded a model of fibroblast mechano-chemo signal transduction capable of accurately predicting fibrosis-related protein expression in response to canonical biochemical factors and mechanical tension. We examined the extent and dynamics of mechano-chemo interactions by simulating biochemical dose-response behavior under varying levels of mechanical stimulation, finding that tension amplified, dampened, or reversed fibroblast responses to individual biochemical cues. Comprehensive simulations of fibroblast responses to over 23,000 combinatory drugs in low- or high-tension contexts identified several drug combinations that adapted fibrotic activity to the local mechanical state, demonstrating the potential of this model to act as a screening tool for targeted therapeutics.

## Results

### Development of Fibroblast Mechanotransduction Network

To develop a fibroblast signaling model capable of sensing and transducing dynamic tension, we performed a manual literature search of intracellular signaling reactions related to fibroblast mechanotransduction and integrated our search with an existing model of fibroblast signaling (Zeigler et al., 2016). Our extended model includes 9 cytokine inputs known to mediate fibroblast activation post-MI as well as mechanical tension, which is transduced via multiple mechanosensors including integrins, stretch-activated ion channels, and the G-coupled protein receptor angiotensin type I receptor (AT1R) (Fig. 1, see Fig. S1 for nodes added to the previous network). We also added several outputs that have been recently shown to mediate myocardial remodeling, including proMMPs 3, 8, and 12, tenascin-c (TNC), osteopontin, and thrombospondin-4 (Frolova et al., 2012; Iyer et al., 2015; Singh et al., 2010; Song et al., 2017), as well as autocrine feedback mechanisms found to regulate fibroblast activation (Ashizawa et al., 1996; Midwood and Schwarzbauer, 2002; Zent and Guo, 2018). The total network, consisting of 109 nodes and 174 reactions, was implemented as a logic-based ordinary differential equation (ODE) model using the open-source Netflux package as previously described (Kraeutler et al., 2010) in which node activity levels and reaction weights are normalized values between 0 and 1 (e.g. an activity of 0.5 represents 50% of maximum activity). All model species, reaction logic, default reaction parameters, and supporting references can be found in Supplemental File S1.

**Figure 1.**
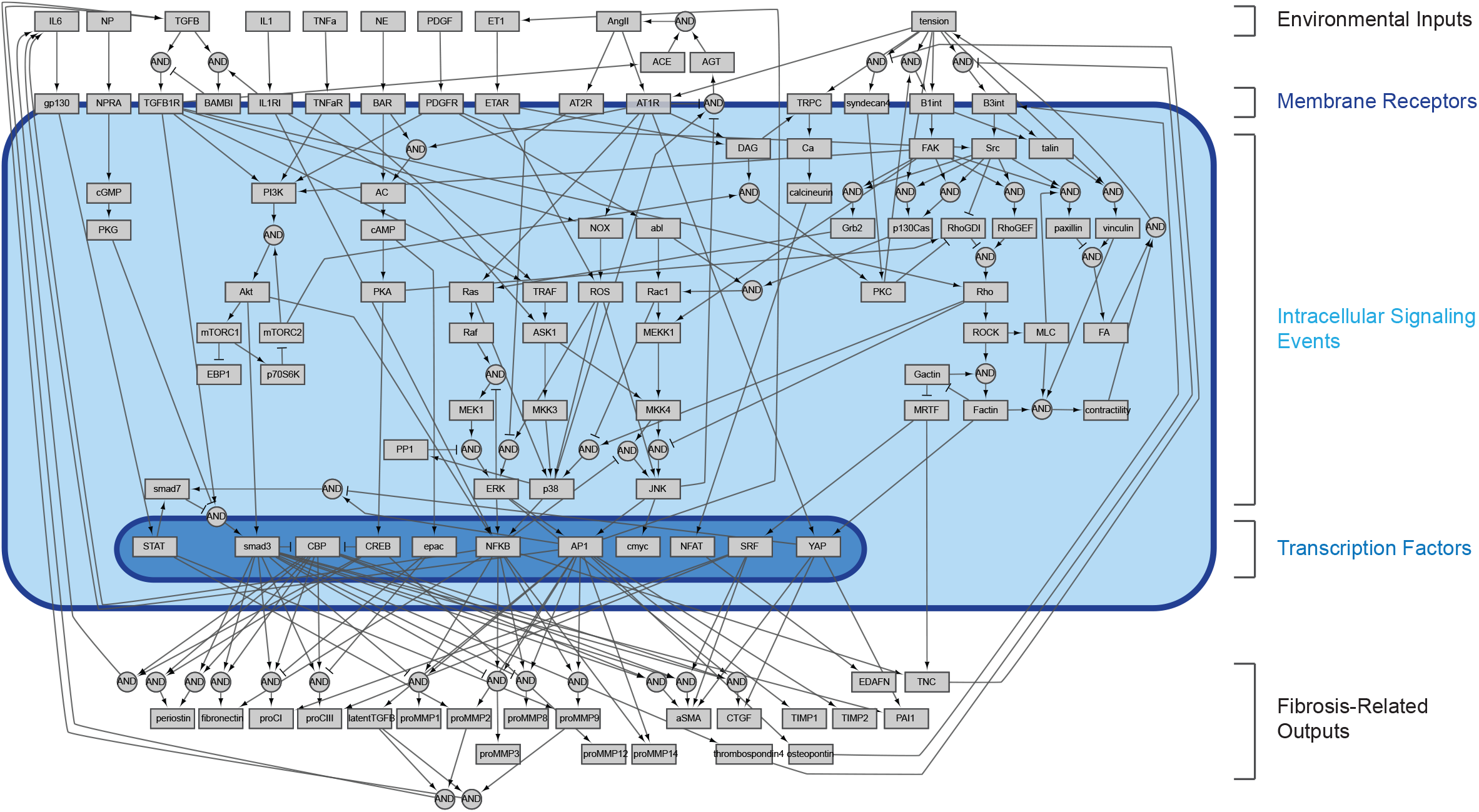
Schematic of expanded logic-based ordinary differential equation model of cardiac fibroblast chemo-/mechano-transduction. Extracellular stimuli, intracellular signaling species, transcription factors, and fibroblast-secreted outputs are represented as nodes (boxes, 109 total). Directed edges represent activating and inhibiting reactions (arrows and T junctions respectively, 162 total) with AND logic indicated by circular nodes.

Model predictions were validated against an independent set of experimental studies by comparing qualitative changes (i.e., increase, decrease, or no change) in node activation to changes in protein activity measured *in vitro*. Parameter sweeps for half-maximal effective concentrations (EC50) and Hill coefficients (n) identified default reaction parameters for optimizing predictive accuracy (Fig. S2, see STAR methods, Model Validation section for full description). For fibroblast-secreted outputs, simulated responses to biochemical and mechanical stimuli correctly predicted qualitative changes in protein secretion in 81.8% (63/77) of simulations (Fig. 2A). Additionally, the model correctly predicted qualitative changes in activation levels of canonical signaling intermediates for 80.5% (33/41) of simulations compared to experimental studies (Fig. 2B). Consistent with previous studies, the model predicted increased expression of collagens, matricellular proteins, and protease inhibitors with exposure to canonical agonists TGFβ1, AngII, platelet-derived growth factor (PDGF), tumor necrosis factor-α (TNFα), and mechanical tension, and secretion of these pro-fibrotic species was accompanied by increased MMP production. Disagreement between model predictions and experimental studies were typically found for studies that reported either decreases or no change in output expression with interleukin-1β (IL1), interleukin-6 (IL6), or NE simulation. However, as described in the next section, these incorrect model predictions could be rectified with the experimentally observed decreases under other input contexts.

**Figure 2.**
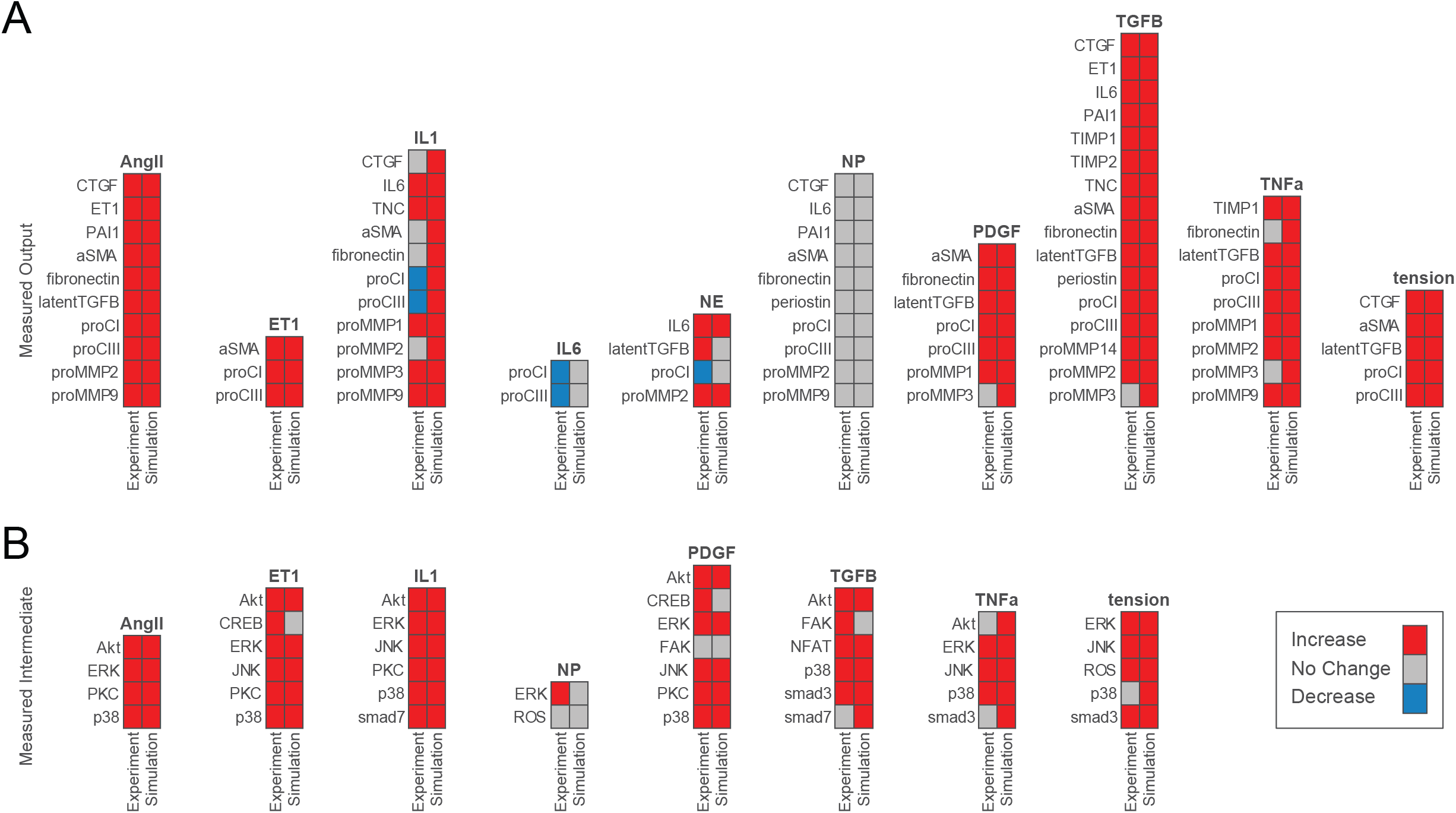
Model accurately predicts qualitative changes in output expression and intermediate activity as measured by independent *in vitro* studies. (A) Expression of cell-secreted outputs and phenotypic markers (i.e. αSMA) were predicted in response to single biochemical stimuli and mechanical tension and compared to an independent set of experimental studies found in literature (46 papers total). (B) Activation of intracellular intermediates representing major signaling pathways were predicted in response to single stimuli and compared to independent experimental studies measuring changes in activity (e.g. by phosphorylation, 27 papers total). All model predictions were categorized based on a ±5% change in activity levels compared to baseline conditions.

### Mechanical Tension Amplifies, Dampens, or Reverses Network Responses to Biochemical Stimuli

Several previous experimental studies have established that combined biochemical and biomechanical signals can interact to produce context-dependent responses from fibroblasts (Bishop et al., 1998; Kural and Billiar, 2016; Lindahl et al., 2002; Merryman et al., 2007; Speight et al., 2016). But the full scope and mechanisms behind these interactions are mostly unclear. We investigated interactions between tension and individual biochemical stimuli computationally by simulating dose-response relationships between biochemical inputs and all model nodes under increasing levels of tension, using area under the curve measurements (AUC) as a metric for total change in activity. We characterized interactions based on changes in each AUC from basal tension to increased levels of tension, categorizing nodes with increased AUC levels with tension as being *amplified*, nodes with decreased AUC levels as being *dampened*, and nodes that changed the direction of activation with tension (i.e. from net activation to net inhibition or vice versa) as being *reversed*.

We found that these interactions were dependent on both the individual inputs as well as the level of tension. Of the 9 biochemical model inputs, AngII, TGFβ1, IL1, TNFα, PDGF, and ET1 all demonstrated sizable changes in dose-response behavior with various levels of tension, based on a 5% change in AUC levels. Using this threshold, a tension level of 0.5 (i.e. 50% of maximum stimulation) amplified, dampened, or reversed an average 56.7% of node AUCs towards these six inputs compared to basal levels of tension (Fig. 3A), and a tension level of 0.9 (i.e. 90% of maximum stimulation) sizably altered an average 60.4% of node AUCs (Fig. 3B). We additionally compared network-wide changes in AUCs between levels of tension as probability distributions, finding that tension significantly altered distributions of dose-response behavior of these six inputs at tension levels of both 0.5 and 0.9 (Fig. S3A, see Table S1 for results of statistical tests). Conversely, tension altered the dose-response behavior of very few nodes with IL6, NE, and NP stimuli, with only 7.03% of node AUCs on average changing sizably towards these inputs for a tension level of 0.9 (Fig. 3B). We found that although tension levels of 0.5 and 0.9 significantly altered probability distributions of changes in AUC for these inputs (Table S1), the relatively small population of nodes with sizable changes suggests that these inputs were largely unchanged with tension., In support of their insensitivity to tension, we found that IL6, NE, and NP all displayed lower degrees of network connectivity compared to the remaining inputs as evidenced by lower measures of closeness centrality and higher measures of average shortest path length and eccentricity (Fig. S3B). While our previous work has shown that these measures do not correlate with functional measures such as sensitivity studies (Zeigler et al., 2016), these topological trends point to a less-influential role of tension for inputs that are relatively less connected to mechanotransduction pathways in the network.

**Figure 3.**
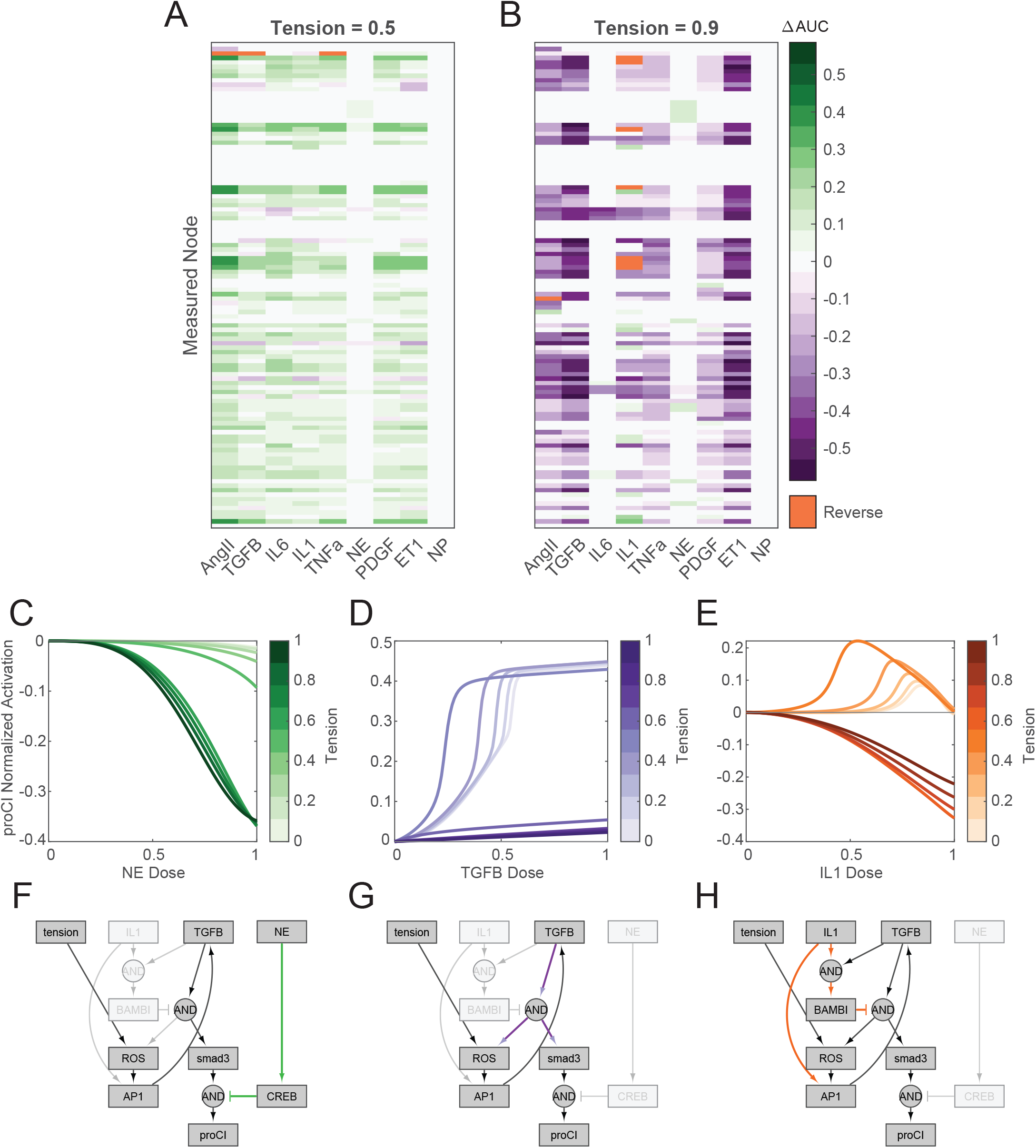
Network-wide responses to biochemical stimuli are dependent on the tensional context. (A-B) Dose-response curves were simulated under elevated levels of tension with incremental doses of each biochemical input, and changes in area under normalized dose-response curves (ΔAUC) were calculated for all nodes in the model in comparison to basal dose-response simulations (i.e. tension = 0.1). Reversal cases were identified based on opposing AUC signs between basal and elevated tension simulations, i.e. nodes that exhibited a positive AUC with basal tension and a negative AUC with elevated tension, and vice versa. (C-E) Examples of mechano-chemo interaction categories identified from network-wide analysis in (A) and (B). Normalized dose-response curves for procollagen I (proCI) expression are shown following simulations with incremental doses of NE (A), TGFβ1 (B), and IL1 (C) under static levels of tension. (F-H) Sub-networks regulating the mechano-chemo interactions shown in (C-E) were determined by manual examination of node activity levels following dose-response simulations using NE (F), TGFβ1 (G), and IL1 (H) stimuli under basal and elevated levels of tension.

Within inputs that demonstrated tension-modulated dose-response behavior, the level of tension applied mediated different types of interactions. Moderate tension levels (reaction weights 0.2-0.5) largely amplified the effects of inputs; of the six inputs demonstrating tension-mediated changes in dose-response behavior, 51.8% of node AUCs were amplified at a tension level of 0.5 on average while 4.43% and 0.46% of node AUCs were dampened or reversed on average, respectively (Fig. 3A). High levels of tension (reaction weights 0.6-0.9) largely dampened network responses to these inputs, with 56.4% of nodes on average decreasing AUCs with the same six inputs compared to 2.75% and 1.22% of node AUCs showing sizable amplification or reversal, respectively (Fig. 3B). Repeated analyses with incremental levels of tension suggested that this change in tension-mediated behavior occurred at a tension reaction weight of 0.6, with an average 40.3±12.1 nodes changing from amplification to dampening at this level for these inputs compared to a reaction weight of 0.5 (Fig. S3A). These changes in sensitivity were indicative of network interactions between tension and biochemical stimuli; we observed that NE-mediated inhibition of procollagen I expression was amplified by tension (Fig. 3C), which can be explained by the logical requirement for tension-induced smad3 activation for effective inhibition by cAMP response element-binding protein (CREB) (Fig. 3D). We also found that tension-mediated desensitization of procollagen I expression towards TGFβ1 can be explained mechanistically by a common autocrine feedback loop integrating both tension and TGFβ1 signals via a reactive oxygen species (ROS)-activator protein 1 (AP1) signaling axis (Fig. 3E-F).

Notably, 4.40% of nodes reversed the direction of dose-response curves with IL1 stimulation with a tension reaction weight of 0.6 or greater compared to baseline tension levels, including intermediate nodes involved in TGFβ1 signaling (TGFβ1R, NOX, ROS), mitogen-activated protein kinase (MAPK) signaling (TRAF, ASK1, MKK3, MKK4, ERK) as well as the expression of procollagen I (Fig. 3G). By comparing these reversed nodes to the network topology, we found that IL1 exerted both stimulatory effects via activation of AP1 and suppressive effects via activation of BMP and activin bound inhibitor (BAMBI), with suppressive behavior dominating procollagen I expression at high IL1 levels (Fig. 3H). While low tension levels allowed for IL1-mediated activation of AP1 and downstream procollagen expression, high levels of tension alone were sufficient to saturate AP1 levels, eliminating the stimulatory effects of IL1 and causing high cytokine doses to inhibit expression. Our model predicts a high degree of interaction between tension and biochemical stimuli, suggesting that the effects of current and potential therapeutics for cardiac fibrosis may be substantially altered by local tension.

### Network Perturbation Analysis Reveals Differential Roles of Tension

While targeted experimental studies of fibroblast signaling have identified numerous pathways mediating extracellular matrix turnover, it is still unclear whether certain mechanisms influence cell behavior more than others and whether these influences are altered with changes in the mechanical context. Using our systems-level model, we calculated changes in network activity following comprehensive knockdowns of individual nodes to functionally assess network-wide effects of perturbations. We calculated each node’s “*knockdown influence”* as the summed magnitudes of all changes in other nodes’ activities upon knockdown of that particular node, and we calculated each node’s “*knockdown sensitivity”* as the summed magnitudes of all changes in that particular node’s activity with knockdown of all other nodes (see STAR Methods, Network Perturbation Analysis section for full description). We repeated simulations for low, medium, and high levels of tension (reaction weights 0.25, 0.5, and 0.75 respectively) and selected for the top 10 scoring nodes in both influence and sensitivity, which indicated differentially responsive sets of nodes (Fig. 4A-C, see Fig. S4 for network-wide sensitivity results).

**Figure 4.**
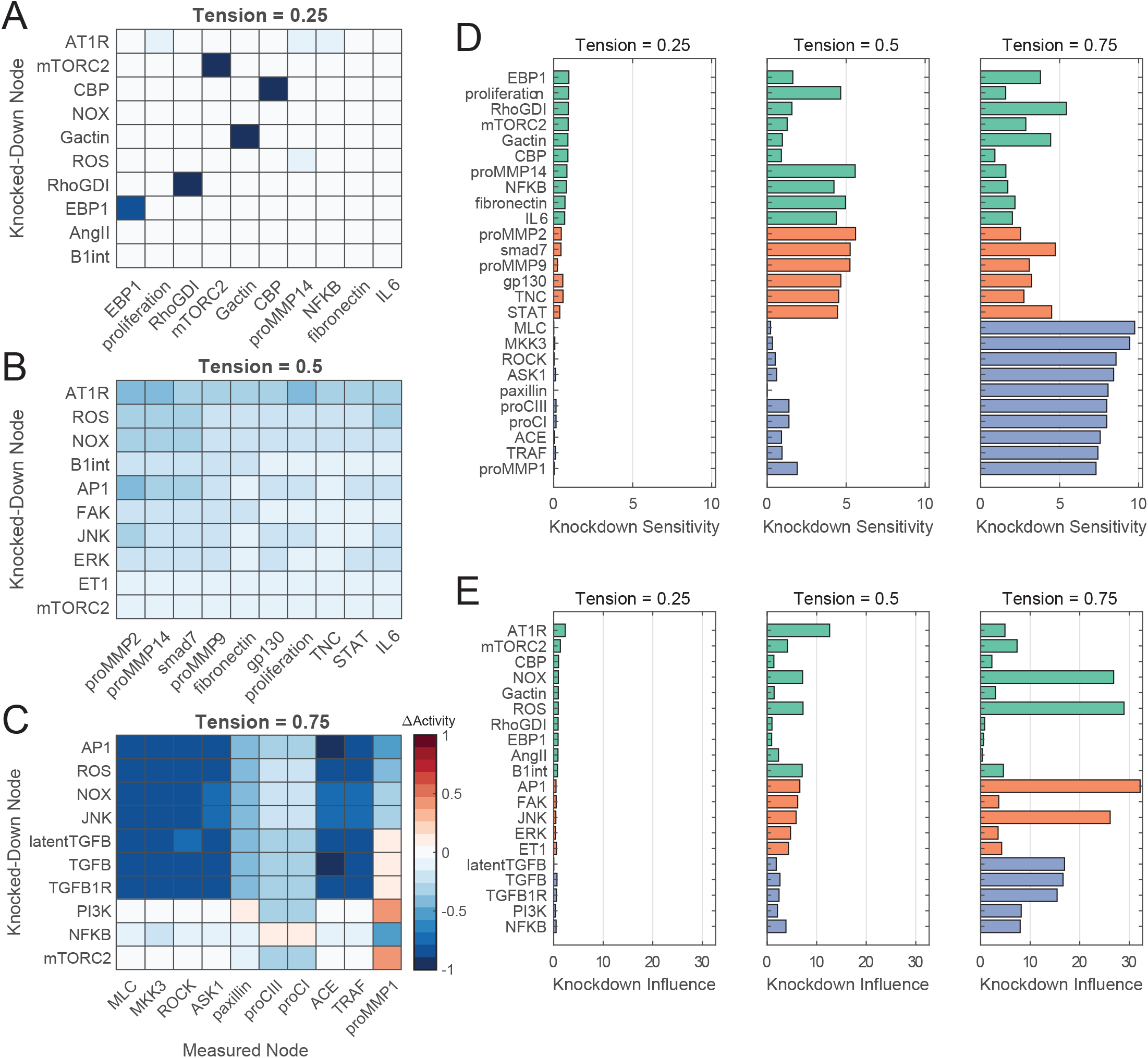
Network perturbation analysis reveals influential pathways of fibroblast mechanotransduction. (A-C) Changes in activity levels for top 10 sensitive nodes with simulated knockdown of top 10 influential nodes at low (A), medium (B), and high (C) levels of tension. Nodes were ranked based on knockdown influence (rows) and knockdown sensitivity metrics (columns), and changes in activity levels with perturbations are compared un-perturbed (i.e. no knockdown) conditions at each tension level. (D-E) Comparison of top-ranked sensitive and influential nodes between levels of tension. Nodes shown reflect all unique nodes that ranked in the top 10 based on knockdown sensitivity (D) or knockdown influence (E) between all levels of tension simulated, and colors reflect nodes that were top-ranked starting at low tension (green), medium tension (orange), and high tension (purple).

At low levels of tension, knockdowns produced only small changes in node activities and correspondingly low influence and sensitivity values (Fig. 4A, D, E). This is not surprising given all node activities are relatively low at this close-to-baseline context. In contrast, medium and high levels of tension demonstrated marked activity changes with knockdown perturbations and exhibited common influencers and unique sets of sensitive nodes. Under medium tension levels, knockdown of AT1R exhibited relatively broad inhibition compared to other top influencers, decreasing activity levels of several proMMPs, fibronectin, TNC, and members of the IL6 pathway by an average of 35.9±5.4% (Fig. 4B). While other top influencers mediated smaller reductions in activity across the top 10 sensitive nodes, NADPH oxidase (NOX), ROS, AP1, and c-Jun N-terminal kinases (JNK) demonstrated an increased capacity to mediate proMMP2, proMMP14, and smad7 levels under medium tension. Under high tension conditions, however, these four nodes ranked as the highest influencers network-wide, mediating decreases in activity for members of the MAPK pathways (80.6±4.3% reduction in activity on average), actin cytoskeleton-related signaling (68.7±22.2% average reduction), and outputs procollagen I, procollagen III, and proMMP1 (30.9±8.9% average reduction) (Fig. 4C). Several members of the TGFβ1 pathway demonstrated similar trends in regulation including latent TGFβ1, suggesting that autocrine feedback may play a role in regulation at high tension.

Rankings of knockdown influence and knockdown sensitivity across all tension conditions suggest that influential mechanisms and their effects were dependent on the tensional context (Fig. 4D-E). While ROS, AP1, JNK, and TGFβ1-related node knockdowns displayed an overall lower degree of influence on the network compared to knockdown of AT1R for low- and medium-tension conditions, knockdown of these nodes under high levels of tension mediated large changes in network-wide activity with knockdown influence levels increasing by up to 4.9-fold compared to medium tension (Fig. 4D). Similarly, different sets of nodes were more sensitive towards perturbations between tension levels, with lower knockdown sensitivity levels of proMMPs 2, 9, and 14 and higher levels of proCI, proCIII, and proMMP1 at high tension relative to medium tension (Fig. 4E). These behaviors suggest that tension acts via different pathways at different levels of stimulation, and cell behavior in a dynamic mechanical environment may not be dictated by a single mechanism alone.

### Fibroblast Signaling Network Predicts Behavior of Current Post-MI Therapeutics

Current post-MI therapeutics aim to reduce cardiac hypertrophy via inhibition or activation of neurohormonal signaling (Ponikowski et al., 2016), both of which also alter cardiac fibroblast behavior and extracellular matrix turnover as a side effect. To test the hypothesis that our model can reproduce experimentally observed effects of current post-MI drugs on fibroblast behavior, we simulated the effects of three drug classes by modulating the maximum activation parameters of individual nodes: (1) angiotensin receptor blockers (ARB) were simulated via knockdown of the AT1R node, (2) neprilysin inhibitors (NEPi), which increase the bioavailability of natriuretic peptides, were simulated via overexpression of the NP node, and (3) ARB-NEPi combination drugs were simulated via combination of (1) and (2) above (see STAR Methods, Drug Effect Comparisons section for full description and Table 1 for simulation parameters).

**Table 1.** Model parameters used for drug effect simulations. Minimum/maximum AT1R inhibitor (AT1Ri) values represent modifiers subtracted from the Y_max_ parameter of the AT1R node, and minimum/maximum neprilysin inhibitor (NEPi) values represent modifiers added to the Y_max_ parameter of the NP node. Non-specified input levels were set to a default value of 0.1. Int.: interpolated from measurements of post-MI cytokine levels *in vivo*.

| Study | TGFB<br>(w) | AngII<br>(w) | NP<br>(w) | Tension<br>(w) | AT1Ri<br>Minimum | AT1Ri<br>Maximum | NEPi<br>Minimum | NEPi<br>Maximum |
| --- | --- | --- | --- | --- | --- | --- | --- | --- |
| (von Lueder et al., 2015) | - | 0.4 | - | 0.3 | 0.01 | 0.5 | 0.05 | 4 |
| (Burke et al., 2019) | 0.4 | 0.4 | 0.6 | 0.3 | 0.5 | 0.5 | 4 | 4 |
| (Ramirez et al., 2014) | Int. | Int. | Int. | 0.1/0.6 | 0.5 | 0.5 | - | - |

We tested whether our model could discern both individual and synergistic effects of post-MI drugs by simulating dose-response relationships between the three drugs above and procollagen I expression. Von Lueder and colleagues previously found that valsartan primarily attenuated collagen expression in cardiac fibroblasts following exposure to AngII, with neprilysin inhibitor LBQ657 acting in a synergistic manner with valsartan but not altering collagen expression alone (von Lueder et al., 2015). Our model independently confirmed this conclusion, as increasing levels of AT1R knockdown combined with NP overexpression further attenuated procollagen I expression compared to AT1R knockdown alone (Fig. 5A). We additionally observed that NP overexpression alone did not appreciably alter procollagen I expression while AT1R knockdown suppressed expression to near-control levels, further supporting the authors’ conclusions of the dominant mechanism for combinatory ARB-NEPi drugs (Fig. S5A-B). This inability of NEPi class treatments to independently regulate AngII signaling can be attributed to a lack of overlap between AngII and NP pathways; AngII stimulation causes the activation of canonical and non-canonical TGFβ1 signaling via production of latent TGFβ1, and although NP overexpression inhibited canonical activation of smad3 by TGFβ1R, non-canonical activation and downstream procollagen expression via Akt remained unchanged. Knockdown of AT1R inhibited both pathways of TGFβ1 signaling, thereby reducing procollagen I levels in a dose-dependent manner.

**Figure 5.**
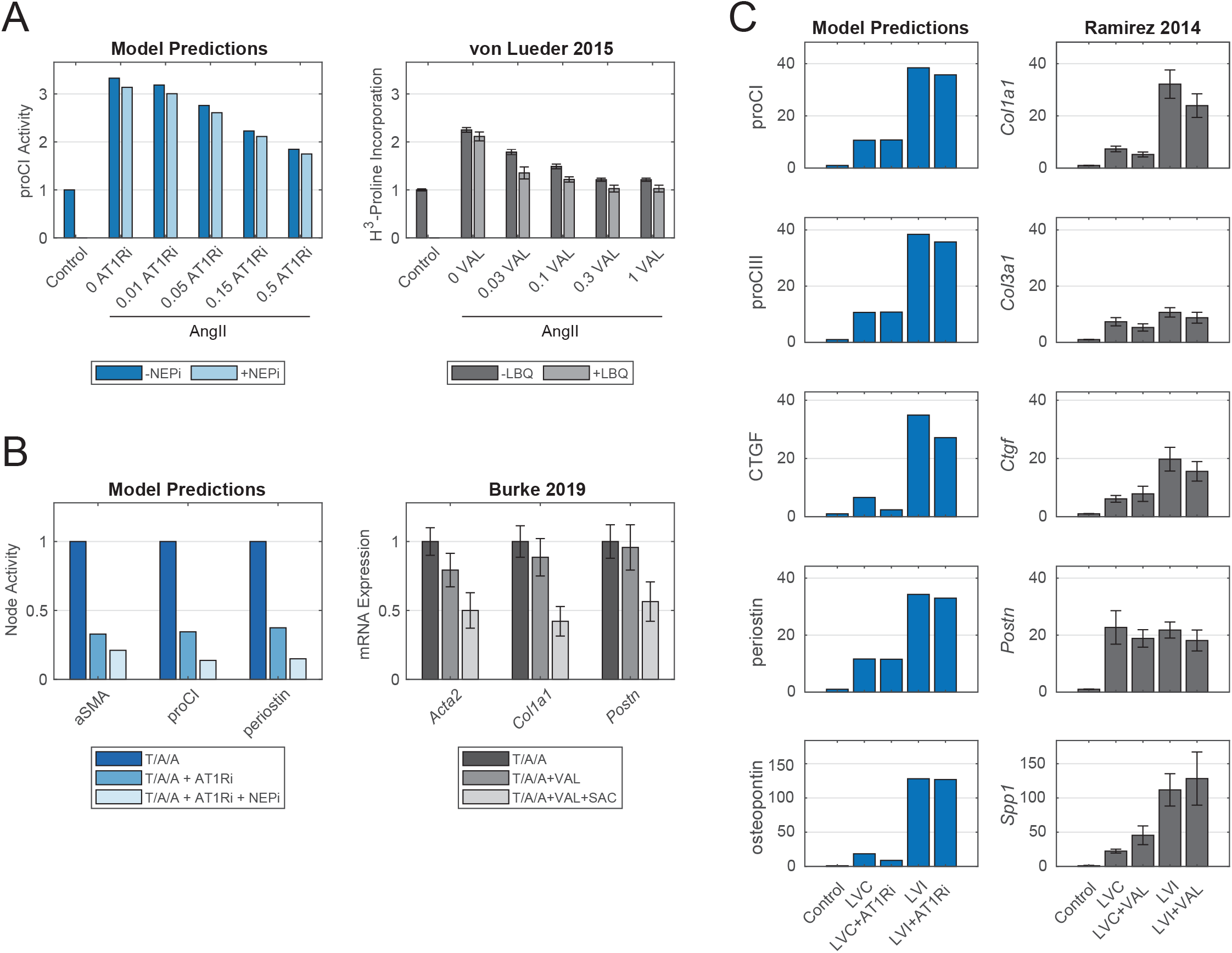
Model-predicted effects of current post-MI drug treatments reflect experimental evidence from independent case studies. (A) Comparison of model-predicted effect of ARB doses with or without NEPi treatment on procollagen I expression with measurements of collagen synthesis in cardiac fibroblasts *in vitro*. Doses for model predictions reflect negative modifiers of the y_max_ parameter of AT1R, doses for the *in vitro* study reflect doses of valsartan in μM, and values reflect fold changes compared to control conditions. (B) Comparison of model-predicted effects of ARB and ARB-NEPi treatments on three output nodes with corresponding gene expression measurements of cardiac fibroblasts *in vitro*. Values reflect fold changes compared to TGFβ1/AngII/atrial NP (T/A/A) conditions. (C) Comparison of model-predicted effects of ARB treatment on five output nodes in representative remote (LVC) and infarct (LVI) zones with corresponding gene expression measurements in post-MI mouse hearts. Values reflect fold changes compared to control (non-infarcted) conditions. All experimental data are represented as mean ± SEM.

We next assessed our model’s capability to predict the expression of multiple gene products and potential mechanisms-of-action following perturbation by the drug classes above. We compared model predictions of three model outputs following ARB or ARB-NEPi treatments to a recent study by Burke and colleagues, who observed synergistic inhibition of genes encoding alpha-smooth muscle actin (αSMA), collagen I, and periostin by combinations of valsartan and the NEPi sacubitril (Burke et al., 2019). We found that model simulations predicted a similar synergistic inhibition of all three gene products in high TGFβ1/AngII/atrial NP (T/A/A) conditions, and although simulations of ARB treatment alone reduced output expression as concluded in the von Lueder study, combinatory treatment in this context further reduced all three model outputs to an average 16.6% of positive controls (Fig. 5B). The authors additionally suggested that sacubitril mediated on fibroblast behavior via activation of PKG and inhibition of the small GTPase Rho, with patient-derived fibroblasts responding to sacubitril or valsartan-sacubitril treatment with decreased fractions of active Rho (Burke et al., 2019). Upon incorporating this interaction into the model reaction logic, model-predicted activation levels of PKG and Rho following NEPi and ARB-NEPi treatments confirm this mechanism-of-action by NP-related treatments, with ARB-only treatments failing to change activity levels of both PKG and Rho compared to TGFβ1/AngII/NP controls (Fig. S5C-D). These comparisons demonstrate the ability of the model topology to predict both output-level changes with drug treatments as well as identify mechanisms-of-action compared to experimental data.

By incorporating known mechanotransduction pathways into network topology, we expanded our model’s capability to predict responses to drug treatments in local cardiac tissue based on differences in tension. This capability is particularly important in the context of an infarcted left ventricle wherein fibroblasts in the infarct scar are subjected to heightened tensile stretches compared to the fibroblasts in the remote, non-infarcted myocardium that are subjected to normal myocardial tensions (Torres et al., 2018). We investigated this capability by simulating the expression of model outputs in response to valsartan treatment in both low- and high-tension contexts, while setting the model cytokine inputs to experimentally matched post-infarct levels as previously described (Zeigler et al., 2020). We compared model output expressions to a study by Ramirez and colleagues, who observed significant differences in fibrosis-related gene expression between infarct and remote zones and negligible effects of valsartan treatment alone using a mouse model (Ramirez et al., 2014). Model predictions generally agreed with their findings, as the degree of tension sizably altered the expression of procollagens, matricellular proteins, and connective tissue growth factor (CTGF) relative to changes in expression with valsartan treatment (Fig. 5C). While we observed variable degrees of error between simulated levels of individual outputs and measured gene expression, our model accurately predicted qualitative differences in drug response based on regional tension.

### Computational Drug Screen Predicts Candidates for Mechano-Adaptive Therapy

The infarcted ventricle presents a spatially dependent need to maintain extracellular matrix content at the infarct region while limiting scar formation in remote regions. While targeted experimental studies involving individual or combinatory post-MI drugs demonstrate efficacy either for individual cell types or bulk tissues, there is currently no evidence of current or potential drugs showing variable effects based on regional mechanics. Our analysis of mechano-chemo interactions above suggests that regional mechanics can alter the effects of positive and negative perturbations, so we investigated whether simulated drug treatments for low- and high-tension conditions could identify drug targets that provide regional therapeutic effects post-MI. Specifically, we sought potential drug targets that adapt matrix-related expression to the regional tension by increasing pro-matrix expression in high tension and decreasing expression in low tension. To that end, we performed a comprehensive screen of individual and combinatory drug targets involving single node knockdown and overexpression and dual node knockdown, overexpression, and knockdown-overexpression combinations for a total of 23,762 drug treatments.

We used a rank-based scoring strategy to score drug treatments based on changes in output expression compared to non-perturbed conditions, calculating a matrix content metric from individual output nodes and ranking treatments by the capacity to decrease or increase this metric with low or high tension stimuli respectively (see STAR Methods, Mechano-Adaptive Drug Screen section for full description). While no individual drug target perturbation achieved these opposing responses (Fig. S6), we found that 450 of 23,544 combinatory perturbations qualitatively altered matrix content based on regional tension, and we identified a cluster of 13 perturbations that scored highly in adapting expression of fibrosis-related outputs to regional tension (Fig. 6A, boxed combinations). This cluster primarily involved dual knockdown of nodes related to the IL6 and Akt/mTOR signaling pathways, with 8 of 13 treatments targeting members of these pathways (Fig. 6B-C). While most of these combinations exerted relatively few changes in individual output levels compared to un-perturbed conditions (Fig. 6B), these combinations primarily acted to increase expression levels of procollagens, periostin, and PAI1 while decreasing proMMPs 1, 3, and 8 with high tension, thus aiding scar formation in high-tension conditions only. Additionally, two treatments showed evidence of increasing proMMP levels and decreasing periostin levels with low tension (Fig. 6B, left-most columns), suggesting that inhibition of the IL6 and Akt/mTOR pathways may simultaneously reduce pro-fibrotic matrix expression in remote tissues with low levels of tension.

**Figure 6.**
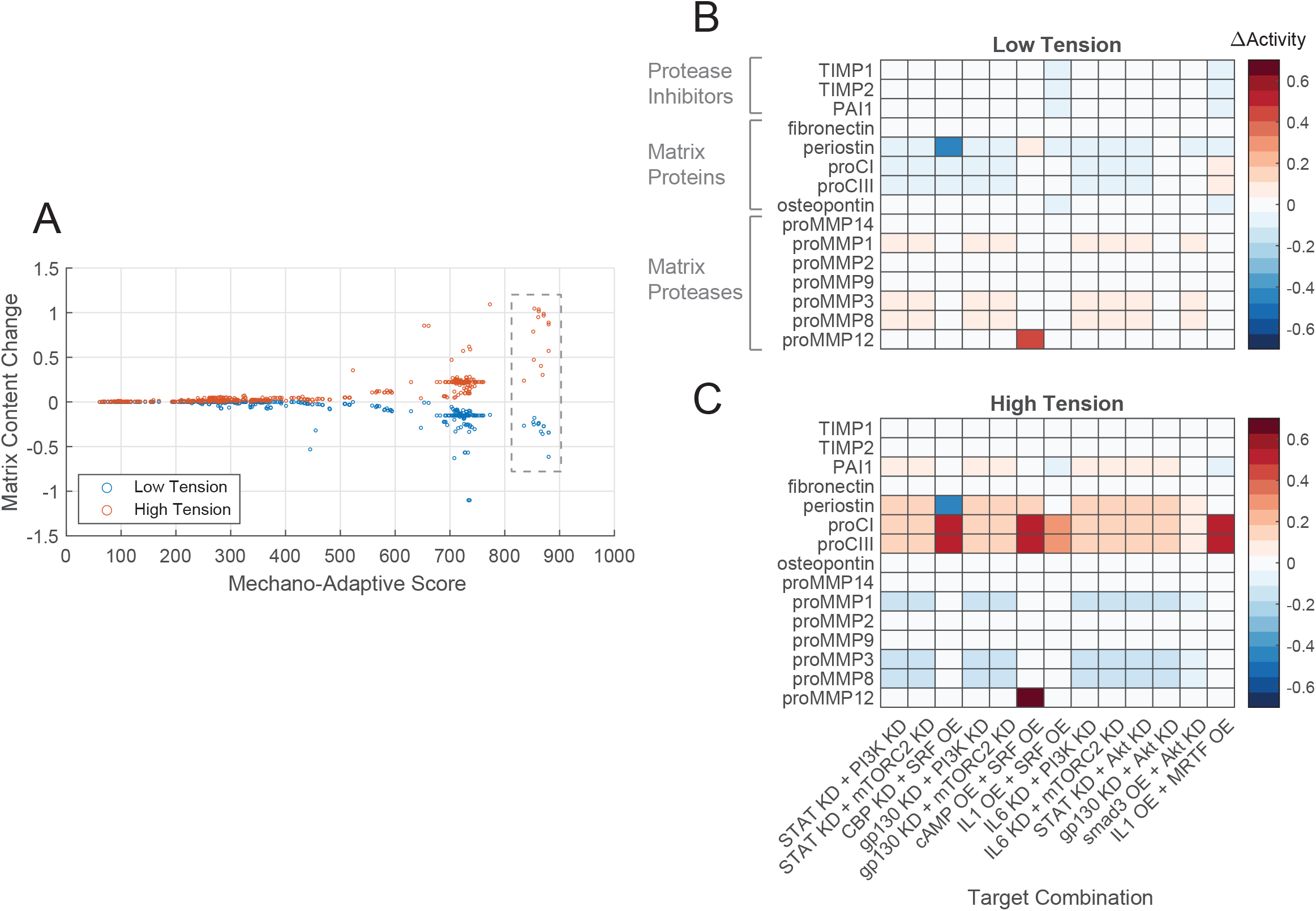
Comprehensive drug screen identifies mechano-adaptive candidates for post-MI fibrosis. (A) Target combinations meeting preliminary threshold for mechano-adaptive behavior (i.e. promoted negative change in matrix content in low tension and positive change in matrix content in high tension) were scored based on rankings of matrix content changes in low- and high-tension simulations. Boxed values represent scores of 13 candidates examined in panels (B) and (C). (B-C) Changes in output expression predicted with mechano-adaptive candidates under tensional contexts, expressed as changes in node activity compared to non-perturbed conditions for each level of tension. Bracketed groups in panel (B) represent categories used for calculations of matrix content in (A).

## Discussion

Spatial control of myocardial fibrosis after MI has remained a challenge due in part to simultaneous signaling from biochemical cues and regional mechanics, both of which mediate fibroblast activation and synthesis of extracellular matrix components (Davis and Molkentin, 2014). In the current study, we employed a computational model of fibroblast signaling to predict the regional expression of fibrosis-related proteins by incorporating known signaling pathways related to canonical biochemical agonists and mechanical tension. We found that the model correctly predicted qualitative changes in fibrotic protein expression with a similar degree of accuracy compared to other logic-based ODE networks (Cao et al., 2020; Wang et al., 2020), and model simulations of current therapeutics such as ARB and ARB-NEPi class drugs accurately reproduced trends in output gene expression observed in cardiac fibroblasts and a post-MI mouse model. Functional analyses identified influential mechano-chemo interactions and preferential pathways of mechanotransduction such as the production of latent TGFβ1 via a NOX-ROS-AP1 signaling axis. We additionally used the model as a targeted screening tool by simulating fibroblast responses to drug combinations in regions of low and high tension, and we identified 13 combinations that adapted the expression of matrix-related proteins to the regional mechanical environment. Our results suggest that this computational approach can aid the development of effective therapies for pathological tissue remodeling which involves simultaneous changes to paracrine signaling and tissue mechanics, and model predictions can generate targeted hypotheses for controlling fibroblast behavior for further experimental investigation.

### Crosstalk Between Biochemical and Biomechanical Factors in Fibroblast Signaling

Increased circumferential stretch in the post-MI infarct zone has been established as both a direct regulator of fibroblast activation and an indirect regulator of biochemical signaling pathways via crosstalk mechanisms (Herum et al., 2017a; Margadant and Sonnenberg, 2010). Experimentally assessing the system-wide extent and dynamics of these interactions would be exceedingly challenging due to time and resource constraints; however, biological network models can overcome this challenge by comprehensively simulating biochemical dose-response relationships under a variety of mechanical contexts. Our findings indicate that tension exerts a wide influence on biochemical signaling in fibroblasts by either amplifying responses to biochemical stimuli in the case of NE, dampening responses in the case of TGFβ1, and even reversing responses in the case of IL1.

The three example cases above share a common connection in tension-mediated expression and activation of latent TGFβ1, an autocrine feedback mechanism shown to influence cardiac fibroblast phenotypes (Hinz, 2015; Lindahl et al., 2002). We found that NE acted to antagonize latent TGFβ1 expression via a cAMP-PKA-CREB signaling axis. Activation of CREB through the use of phosphodiesterase inhibitors has previously been shown to antagonize fibrotic expression of lung and cardiac fibroblasts (Miller et al., 2011; Wójcik-Pszczoła et al., 2020) as well as cardiac fibrosis in a pressure overload mouse model (Gong et al., 2014), suggesting a potential mechano-responsiveness of this mechanism. Simulated fibroblast responses to TGFβ1 indicated a synergistic response under tension with latent TGFβ1 activation and output expression occurring for lower input doses. This trend is consistent with experiments involving valvular myofibroblasts, which exhibited earlier and greater induction of bioactive TGFβ1 and aSMA protein levels when subjected to combined tension/TGFβ1 compared to either stimulus alone (Merryman et al., 2007).

We additionally found that stimulation with IL1 produced opposite effects depending on the level of tension, with IL1 increasing procollagen I expression for low to medium tension levels and decreasing expression for high tension levels through activation of BAMBI and downregulation of downstream targets of TGFβ1R. As a positive regulator, IL1 upregulates MAPK and NFkB signaling pathways via myeloid differentiation factor-88 (MyD88) (Van Tassell et al., 2015), and both IL1R^−/−^ and MyD88^−/−^ mice display reduced fibrosis and progression towards heart failure in a model of acute myocarditis (Blyszczuk et al., 2009). As a negative regulator, IL1 upregulates the expression of BAMBI to act as a decoy receptor for TGFβ1, thereby reducing downstream signaling (Saxena et al., 2013). In BAMBI^−/−^ mice, pressure overload worsened fibrosis compared to wild-type mice in a TGFβ1-dependent manner, suggesting that this negative regulation is dependent on both mechanical stress and TGFβ1 stimulation (Villar et al., 2013). IL1 has additionally been shown to reverse the myofibroblast phenotype in valvular interstitial cells with culture on stiff substrates (Aguado et al., 2019), further suggesting that context-dependent regulation by IL1 may be a determining factor of fibroblast phenotype. Our analysis of mechano-chemo interactions above suggests that computational approaches can reveal context-specific mechanisms of disease progression and reinforces the need to account for various environmental factors (such as local biomechanics) in developing therapeutic strategies.

### Influential Mechanisms Controlling Fibroblast Mechanotransduction

Fibroblasts transduce changes in their local mechanical environment using both well-known mechanosensors such as integrins and newly-discovered mechanisms such as piezo channels and actin-dependent translocation of YAP/TAZ (Herum et al., 2017a). While these mechanisms have all been shown to alter cell phenotypes individually, it is still unclear as to whether individual pathways influence cell behavior over others. Network perturbation analysis predicted that tension is primarily mediated by an autocrine feedback mechanism in which a NOX/ROS/JNK/AP1 signaling axis stimulates latent TGFβ1 expression to drive further NOX activation as well as procollagen expression via smad3. The secondary activation of TGFβ1 has been demonstrated for various stimuli in fibroblasts such as TNFα and AngII (Gao et al., 2009; Voloshenyuk et al., 2011) as well as for other cell types by influencing changes in phenotype related to extracellular matrix deposition and angiogenesis (Hamzeh et al., 2015; Joki et al., 2000; Zheng et al., 2001). Inhibition of NOX2 and NOX4 in AngII-infusion mouse models has additionally been shown to attenuate interstitial collagen deposition, further implicating NOX/ROS signaling as a central influencer of cardiac fibrosis (Johar et al., 2006; Zhao et al., 2015). Our model prediction that this pathway primarily influences fibrosis-related protein expression suggests that perturbation of this pathway may induce larger reductions in cardiac fibrosis compared to others, and experimental studies investigating this behavior could provide a basis for developing targeted anti-fibrotic therapies.

### Fibroblast Network Model as a Screening Tool for Targeted Therapeutics

The heterogenous nature of myocardial infarctions creates a clear need for regionally targeted therapies for tissue remodeling, with the infarct zone requiring scar formation to provide local structural integrity and remote zones requiring scar downregulation to prevent adverse tissue stiffening. Loss of cardiomyocytes additionally presents different mechanical conditions between the infarct and remote zones (Torres et al., 2018), and our model predictions above and experimental evidence suggest that myofibroblast-mediated tissue remodeling is dependent on this mechanical context (Watson et al., 2012; Waxman et al., 2012). Previous trials of anti-fibrotic therapies have demonstrated the need to account for local increases in circumferential stretch in infarcted tissue. Studies inhibiting known mediators of excessive fibrosis such as TGFβ1, osteopontin, and TIMP3 have shown increased risk of left ventricular thinning and cardiac rupture, early complications associated with impaired scar formation at the infarct zone (Hammoud et al., 2011; Ikeuchi et al., 2004; Trueblood et al., 2001) - an indication that therapies designed to inhibit global fibrosis are not sufficient to promote healthy scar formation where it is needed. A further complication of post-infarct fibrotic control is the lack of evidence of regional differences in tissue remodeling for current and potential drugs. While our comparisons of model-predicted and experimentally tested ARB and NEPi treatments above demonstrated the inhibitory effects of these drugs in a single context, not enough evidence exists to determine whether these drugs produce therapeutic effects in both remote and infarcted zones.

We investigated the ability of our model to predict regional effects of drug treatments based on regional mechanical tension, and our computational screen of 23,762 drug treatments yielded 13 candidates that adapted expression of extracellular matrix-related proteins to the regional tension. These candidates implicated the IL6 and Akt/mTOR signaling pathways as well as the IL1/cAMP and MRTF/SRF pathways as mediators of mechano-adaptive behavior. Perturbations involving the IL6 and IL1 inflammatory pathways individually have yielded mixed results clinically; while several animal studies and clinical trials perturbing IL6 and IL1 signaling have concluded that knockdown of these pathways suppresses cardiac fibrosis and mediates gain of function due to MI or factors associated with MI (Abbate et al., 2013; Hilfiker-Kleiner et al., 2010; Ma et al., 2012), others have concluded that IL6 and IL1 knockdown exacerbate fibrosis post-MI or in response to MI-associated biochemical factors (Dziemidowicz et al., 2019; Obana et al., 2010; Turner, 2014). The results from these studies indicate that the influence of these pathways on tissue remodeling is highly context dependent with varying responses between treatment time courses and methods of antagonism, and our drug screen indicates that mechanical signaling additionally modifies cellular and tissue responses to these drugs. While we found no clinical evidence of tissue remodeling in response to the model-predicted drug combinations above, these results suggest that pairing of anti-inflammatory treatments with those for additional signaling pathways may provide an additional advantage in controlling regional cardiac fibrosis, and our predictions provide testable hypotheses for further investigation. Moreover, this model-based approach towards target discovery provides substantial value for potential drug development by focusing efforts on drugs or drug combinations with strong mechanistic evidence of therapeutic efficacy. This model and approach can additionally be expanded to investigate fibrosis across multiple disease states and for individual patient conditions to stratify patients based on predicted responses to current drugs or to identify drug candidates for subpopulations of patients.

### Study Limitations and Future Directions

One major limitation of the current study stems from the need for quantitative predictions of fibroblast signaling processes and matrix production. While the current model was able to make semi-quantitative predictions of cell behavior based on network topology and normalized reaction parameters, data-driven reaction parameters are necessary both to fully investigate mechanisms-of-action of fibrotic activity and to relate signaling processes to experimental measures of fibrosis. Phospho-proteomic datasets generated using methods such as reverse-phase array provide direct analogs to intracellular protein activation or inhibition as well as the scope of measurements required for systems-level modeling, and previous studies have demonstrated the utility of these datasets in inferring cell signaling networks and fitting literature-curated network parameters (Alexopoulos et al., 2010; Osmanbeyoglu et al., 2017; Terfve and Saez-Rodriguez, 2012). Future implementation of *in vitro* or clinical phospho-proteomic data would improve the accuracy of model predictions and reveal additional mechanistic insight into fibroblast behavior in the post-MI environment.

An additional limitation of the current model is that of scope as it relates to both cell signaling processes as well as additional levels of regulation. While the current model was developed based on direct experimental evidence of signaling interactions stemming from selected biochemical and mechanical stimuli, we must acknowledge that additional stimuli and signaling pathways play a role in cardiac fibroblast signaling, such as interferon-γ and epidermal growth factor receptor signaling pathways (Levick and Goldspink, 2014; Liu et al., 2018). While not meeting our requirements for inclusion (see STAR Methods, Fibroblast Signaling Model Development section for criteria), we expect these additional signaling mechanisms to aid in model predictions across the diverse range of stimuli represented *in vivo*. It is also important to note that a variety of biomechanical factors alter fibroblast behavior in the post-infarct environment in addition to circumferential stretch (classified as “mechanical tension” in the current study), such as tissue stiffening, compression, and radial and longitudinal stretch (Clarke et al., 2016). While these applied loads are sensed and transduced via similar mechanisms as circumferential stretch, the type, magnitude, and directionality of applied loads differentially mediate fibroblast behavior and matrix turnover (Herum et al., 2017b; Wang et al., 2004), and thus a more robust understanding of the interactions between biomechanical stimuli is needed. Lastly, much evidence exists to suggest that many additional dimensions of regulation mediate cardiac fibrosis including transcriptional regulation (Lacraz et al., 2017), remodeling of collagen fibers (Richardson et al., 2015), paracrine signaling involving inflammatory cells, immune cells and myocytes (Mouton et al., 2018), and additional modes of cell signaling (e.g. microRNA, matrix degradation products) (Lindsey et al., 2015; Thum et al., 2008). Several groups have made progress in modeling regulation within these levels in the heart (Chen et al., 2019; Rouillard and Holmes, 2012; Skelly et al., 2018), and while incorporating all factors into a model-based approach is a non-trivial effort, a combined, multi-dimensional approach will ultimately be necessary to fully predict tissue remodeling given the range of variables involved in post-infarct wound healing.

Through our simulated studies of fibroblast signaling and fibrotic activity above, we demonstrated the utility of computational models of mechano-chemo signal transduction in identifying mechanisms-of-action governing cell behavior in complex environments. While *in vitro* and *in vivo* evidence validating these results are crucial for translating this mechanistic insight into clinical solutions for cardiac fibrosis, this model-driven approach can be advantageous in focusing experimental efforts towards promising candidates that limit scar formation to regions where it is beneficial to do so. Excitingly, our simulations identified 13 novel drug target pairs whose modulation resulted in desirable mechano-adaptive effects - namely, an increase in matrix accumulation within the infarct zone (where scar material is beneficial to maintain structural integrity) and a simultaneous decrease in matrix within the remote zone (where scar material is detrimental to myocardial mechanical and electrical function). Future phospho-proteomic studies of fibroblast signaling in combination with targeted studies of influential signaling pathways will further enable the development of targeted treatments for cardiac remodeling post-MI, and further tailoring of this network towards other pathological states will enable similar mechanistic insight and opportunities for therapeutic development for a range of fibrosis-related diseases.

## Acknowledgements

This study was supported by grants from the National Institutes of Health (GM121342, HL144927) and the American Heart Association (17SDG33410658). We would like to thank the Cyberinfrastructure Technology Integration group at Clemson University for generous allotment of compute time on the Palmetto high performance computing resource. We would also like to thank Dr. Jeff Holmes at the University of Alabama - Birmingham and Dr. Jeff Saucerman at the University of Virginia for rich and thoughtful discussion around the ideas motivating this work.

## Author Contributions

Conceptualization: J.D.R and W.J.R; Methodology: J.D.R and W.J.R; Investigation: J.D.R; Data Curation: J.D.R; Writing – Original Draft: J.D.R; Writing – Review & Editing: J.D.R and W.J.R; Funding Acquisition: W.J.R.

## STAR Methods

### Resource Availability

#### Lead Contact

Further information and requests for resources should be directed to and will be fulfilled by the Lead Contact, William Richardson.

#### Materials Availability

This study did not generate new unique reagents.

#### Data and Code Availability

The code generated during this study is available at GitHub (https://github.com/SysMechBioLab/Fibroblast_Signaling_Network_Model). The published article includes all model and validation databases generated or analyzed during this study.

### Method Details

#### Fibroblast Signaling Model Development

Our previously published model of cardiac fibroblast signaling was expanded using a manual literature search of over 100 peer-reviewed studies to include newly identified proteins and/or signaling pathways associated with fibroblast mechanotransduction, as well as several secreted proteins newly-associated with cardiac fibrosis. New individual proteins (model nodes) and signaling reactions (edges) were identified from studies that concluded direct interactions between proteins altering protein activity (e.g. by phosphorylation). New nodes and/or edges were included if at least two independent studies contained evidence of either activation of a node or activation of another species by a node in either cardiac or other fibroblast subtypes. As previously described (Tan et al., 2017; Zeigler et al., 2016), the final network was implemented as logic-based ordinary differential equations in which activity levels of all nodes were modeled as a system of Hill equations, and all activity levels and input stimuli were represented as normalized values between 0 and 1. Logical NOT, AND, and OR gates were used for inhibitory and complex signaling interactions by applying logical operations 1 − *f*(*x*), *f*(*x*)*f*(*y*), and *f*(*x*) + *f*(*y*) − *f*(*x*)*f*(*y*) to Hill equations respectively. The system of differential equations was constructed using the open-source Netflux package for MATLAB (https://github.com/saucermanlab/Netflux), and all subsequent analyses were conducted using MATLAB (Mathworks, Natwick MA). Visualizations of the full network and sub-networks were created using Cytoscape software (Shannon et al., 2003).

### Quantification and Statistical Analysis

#### Model Validation

Qualitative changes in select model outputs and signaling intermediates were compared between model predictions and independent experimental studies. A set of studies independent from model development (47 papers total) were selected based on direct measurement of either output secretion or activity of a signaling intermediate in response to a single input stimulus in fibroblasts. Outputs included in this set were either collagens, matrix proteases, protease inhibitors, matricellular proteins, autocrine signaling species, or αSMA as a marker of fibroblast activation. Intermediates included were limited to members of canonical signaling pathways (i.e. Akt, CREB, ERK, JNK, p38, PKC, smad3, ROS), and only intermediates with direct measurements were included in the final set. Model predictions were performed by simulating basal conditions (all input weights = 0.1) for 80 s, followed by simulating single input stimuli (weight = 0.8) for an additional 240 s in order to allow for the system to reach steady-state activity levels. Because studies used for validation conducted *in vitro* experiments on tissue culture plastic, a tension input weight of 0.4 (40% of maximum stimulation) was used for all validation simulations. Changes in intermediate and output node activity from basal to stimulated conditions were binned into either “increase”, “decrease”, or “no change” categories using ± 5% change in activity levels, which has been shown previously to distinguish qualitative changes in behavior for single-stimulus simulations (Tan et al., 2017; Wang et al., 2020). To fit default model parameters to best describe experimental data, parameter sweeps of Hill coefficient (n) and half-maximal effective concentration (EC50) were conducted to maximize the accuracy of model predictions, using the percentage of correct predictions as a metric. Maximum activation levels (Y_max_) were set to a default value of 1, and time constants (τ) were set to default values of either 0.1, 1, or 10 corresponding with species involved in intracellular signaling reactions, ligand-receptor reactions, or transcription-translation events respectively. Reaction weights (w) were set to default values of 1 with the exception of autocrine feedback-associated reactions, which were set to default values of 0.8 due to the lowered probability of these reactions resulting diffusion of output proteins or sequestering by the extracellular matrix (Hinz, 2015). Values for n and EC50 found to maximize predictive accuracy were used with the other described parameters for all subsequent simulations. All model species, reactions, and assigned parameters are detailed in Supplemental File S1.

#### Mechano-Chemo Interaction Analysis

Dose-response behaviors of biochemical stimuli were simulated for incremental levels of the ‘tension’ input reaction to assess interactions between mechanical and biochemical stimulations. For 0.1 increments in tension reaction weight ranging from 0.1 (10% of maximum stimulation) to 0.9 (90% of maximum stimulation), steady-state activity levels of all nodes in the network were simulated following stimulation of each biochemical input reaction (AngII, TGFβ1, IL6, IL1, TNFα, NE, PDGF, ET1, and NP) for 240 s, ranging in reaction weight from 0 to 1 in 0.01 increments. The resulting dose-response curves were normalized to steady-state activity levels for a biochemical input of 0, and the total response of each node to a biochemical stimulus was calculated as the area under the normalized curve (AUC). Changes in AUC due to tension (ΔAUC) were calculated for each tension level as the difference between AUCs with added tension and with baseline tension (tension = 0.1). Sizeable mechano-chemo interactions were identified using a threshold of ±5% in ΔAUC. ΔAUCs greater than or equal to 5% were categorized as amplifying interactions (tension increases node response to a biochemical stimulus), ΔAUCs less than or equal to −5% were categorized as dampening interactions (tension decreases node response), and AUCs with opposite signs between baseline and tension conditions and exceeded the threshold of ±5% were classified as reversal interactions (tension causes biochemical stimulation to exert the opposite effect on node activity). This threshold has been shown previously to distinguish qualitative changes in activity levels with individual node perturbations (Tan et al., 2017; Wang et al., 2020), and while these studies did not use an integrated AUC metric for comparison, our comparison of AUCs requires greater peak activation levels to meet the threshold, thereby imposing a stricter comparison than these previous studies.

To compare the extent of interactions between tension levels for each input, two-sample Kolmogorov-Smirnov tests were conducted for distributions of ΔAUCs, comparing (1) distributions for tension = 0.2 vs. tension = 0.5; (2) distributions for tension = 0.2 vs. tension = 0.9; and (3) distributions for tension = 0.5 vs. tension = 0.9. Corrections for multiple comparisons were made using the Benjamini-Hochberg procedure (Benjamini and Hochberg, 2009), and adjusted p-values are displayed in Supplemental Table S1. As an additional measure supporting the input-related differences in interactions found above, topological network parameters were computed using the NetworkAnalyzer plugin for Cytoscape (Doncheva et al., 2012).

#### Network Perturbation Analysis

To identify influential signaling mechanisms across various mechanical contexts, a series of node knockdowns were simulated for several levels of tension. For tension reaction weights of 0.25, 0.5, and 0.75, basal conditions were simulated for 80 s (all other input weights = 0.1) followed by knock down of individual nodes (Y_max_ = 0.1) for 240 s. Steady-state activity levels of all nodes were measured across knockdowns of all nodes, and changes in activity (ΔActivity) were calculated as the difference between node activity after knockdown and basal activity. Knockdown sensitivity of each node was calculated as the total change in activity of the node across all knockdown simulations, and knockdown influence of each node was calculated as the total change in activity of all other nodes in the network upon knockdown.

#### Drug Effect Comparisons

Predicted responses of the network to currently prescribed pharmacologic therapies were compared three studies measuring cardiac fibroblast or myocardial tissue gene expression or protein expression *in vitro* or *in vivo* (Burke et al., 2019; von Lueder et al., 2015; Ramirez et al., 2014). All drugs were simulated by knockdown or overexpression of individual nodes: angiotensin receptor blocker valsartan (VAL), simulated by applying a negative modifier to Y_max_ for the AT1R node, neprilysin inhibitors LBQ657 (LBQ) and sacubitril (SAC), simulated by applying a positive modifier Y_max_ for the NP node, and angiotensin receptor neprilysin inhibitors LCZ696 (VAL+LBQ) and sacrbitril/valsartan (VAL+SAC). All negative control simulations were conducted using input reaction weights of 0.1, and all input reaction weights and drug perturbation levels for *in vitro* studies were chosen to maximize the dynamic range of network responses. Input reaction weights for *in vivo* studies were interpolated from experimental data of cytokine levels post-MI as previously described (Zeigler et al., 2020) using a time point of 4 weeks post-infarction, and tension reaction weights for control and infarct zones were set to 0.1 and 0.6, respectively. All input reaction weights and perturbation conditions can be found in Table 1.

#### Mechano-Adaptive Drug Screen

A computational screen of single and double network perturbations was conducted to identify drug targets with potential to adapt fibrotic activity to the local mechanical environment. Biochemical input reaction weights used for all simulations were interpolated using the method described for *in vivo* study comparisons at a time point of 2 weeks post-infarction. To represent remote (low tension) and infarcted (high tension) zones post-MI, tension reaction weights of 0.1 and 0.6 were used, respectively. For each simulation, baseline conditions were simulated using the input reaction weights above for 80 s, and single or double perturbations were simulated by setting Y_max_ values of nodes to 0.1 (knockdown) or 5 (overexpression) for 240 s. Single perturbations were simulated for all individual nodes as both knockdown and overexpression for a total of 218 perturbations, and double perturbations were simulated for all node combinations as double knockdown, double overexpression, or knockdown-overexpression combinations for a total of 23,544 perturbations.

Perturbations that exhibited mechano-adaptive behavior were identified using a rank-based score metric based on changes in activity levels for model outputs (ΔActivity). For each perturbation and tension level, ΔActivity levels were calculated as the difference in each node’s activity between baseline and perturbed conditions, and changes in overall matrix content (MCC) for each perturbation and tension level were calculated based on ΔActivity levels for all procollagens and matricellular proteins (ΔActivity_Matrix_), ΔActivity levels for all MMPs (ΔActivity_MMP_), and ΔActivity levels for all protease inhibitors (ΔActivity_Inhib_): *MCC* = ∑ Δ*Activity_Matrix_* − Δ*Activity_MMP_* + Δ*Activity_Inhib_*. Output nodes were categorized as follows: ΔActivity_Matrix_: proCI, proCIII, fibronectin, periostin, osteopontin; ΔActivity_MMP_: proMMPs 1, 2, 3, 8, 9, 12, 14; ΔActivity_Inhib_: TIMP1, TIMP2, PAI1. To identify perturbations that (1) qualitatively alter matrix content in a desirable manner based on the tensional context and (2) maximize differences in low- and high-tension expression, a two-step procedure was used to score mechano-adaptive behavior from MCC values:

i. Perturbations were first filtered based on the numeric signs of MCC values at each tension level to identify categorically desirable behavior. Perturbations that exhibited negative MCC values (i.e. greater anti-fibrotic activity than pro-fibrotic activity) at low tension and positive MCC values at high tension (i.e. greater pro-fibrotic activity than anti-fibrotic activity) were retained for subsequent steps, and all others were removed.
ii. Retained perturbations were then ranked based on MCC values at each tension level. MCC values at low tension were ranked in ascending order (i.e. low to high values), and MCC values at high tension were ranked in descending order (i.e. high to low). The mechano-adaptive score for each perturbation was then determined by summing each perturbation’s rank for low and high tension and subtracting two times the sum from the number of retained perturbations, thereby denoting highly adaptive perturbations with high scores and less adaptive perturbations with low scores with a minimum score of 0.

## Supplemental Information Titles and Legends

**Supplemental File S1, Related to Figure 1. Fibroblast mechano-chemo signal transduction model.** Database detailing all nodes, reactions, model parameters, and references used for construction of the signaling network.

**Supplemental File S2, Related to Figure 2. Model validation database.** Database detailing all qualitative input-output and input-intermediate relationships and references used for validation of the signaling network with independent experimental studies.

