## Supplemental Figures S1-S6 and Table S1 for "Fibroblast Mechanotransduction Network Predicts Targets for Mechano-Adaptive Infarct Therapies"

**Supplemental Figure 1**

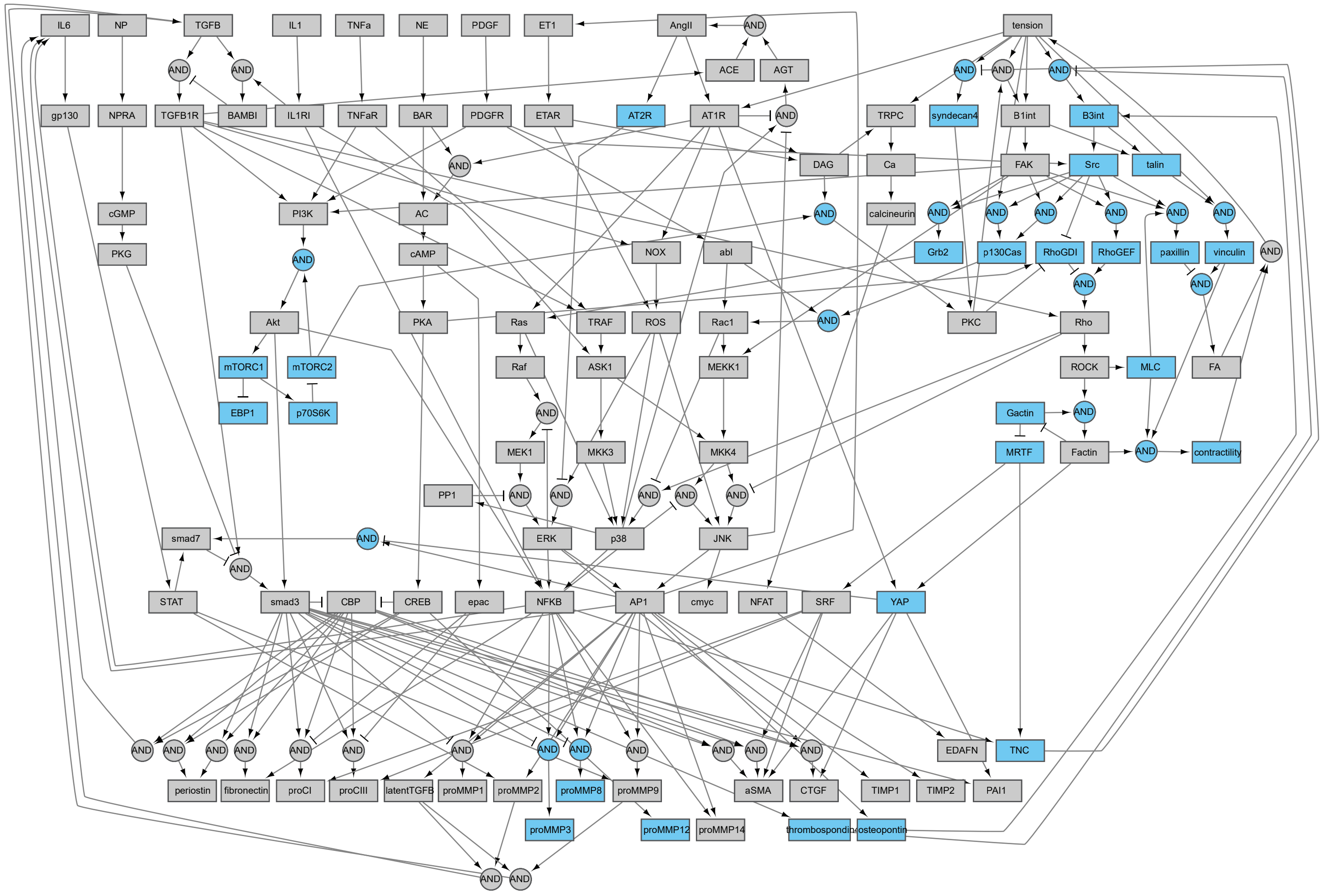


**Figure S1, Related to Figure 1. Schematic of fibroblast chemo-/mechano-transduction network modifications for current study.** Added nodes relating to mechanotransduction mechanisms, mechano-chemo crosstalk mechanisms, and additional secreted outputs are highlighted in blue.

Supplemental Figure 2

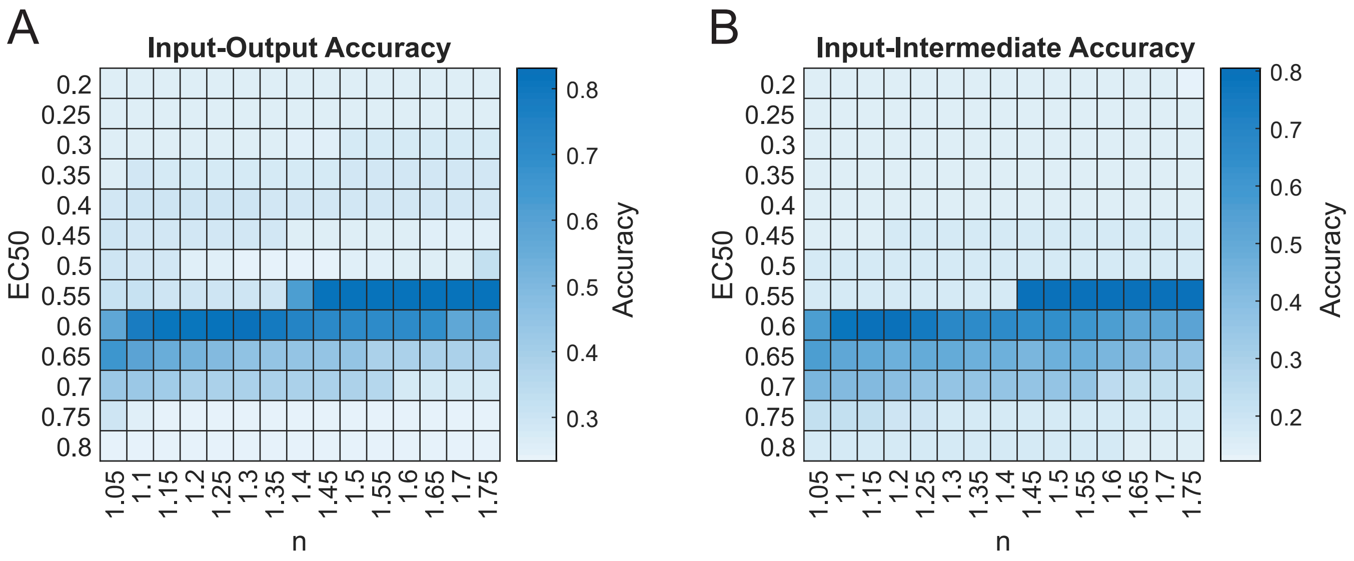


**Figure S2, Related to Figure 2. Optimization of reaction parameters for maximizing accuracy of qualitative predictions.** Parameter sweeps of half-maximal effective concentration (EC50) and Hill coefficients (n) were conducted by simulating changes in output node expression (A) and intermediate node activity (B) in response to single input stimuli (see STAR methods, Model Validation section for full description). Shading of parameter combinations represents the total proportion of correct qualitative predictions compared to each independent experimental validation set used in Figure 2.

### Supplemental Figure 3

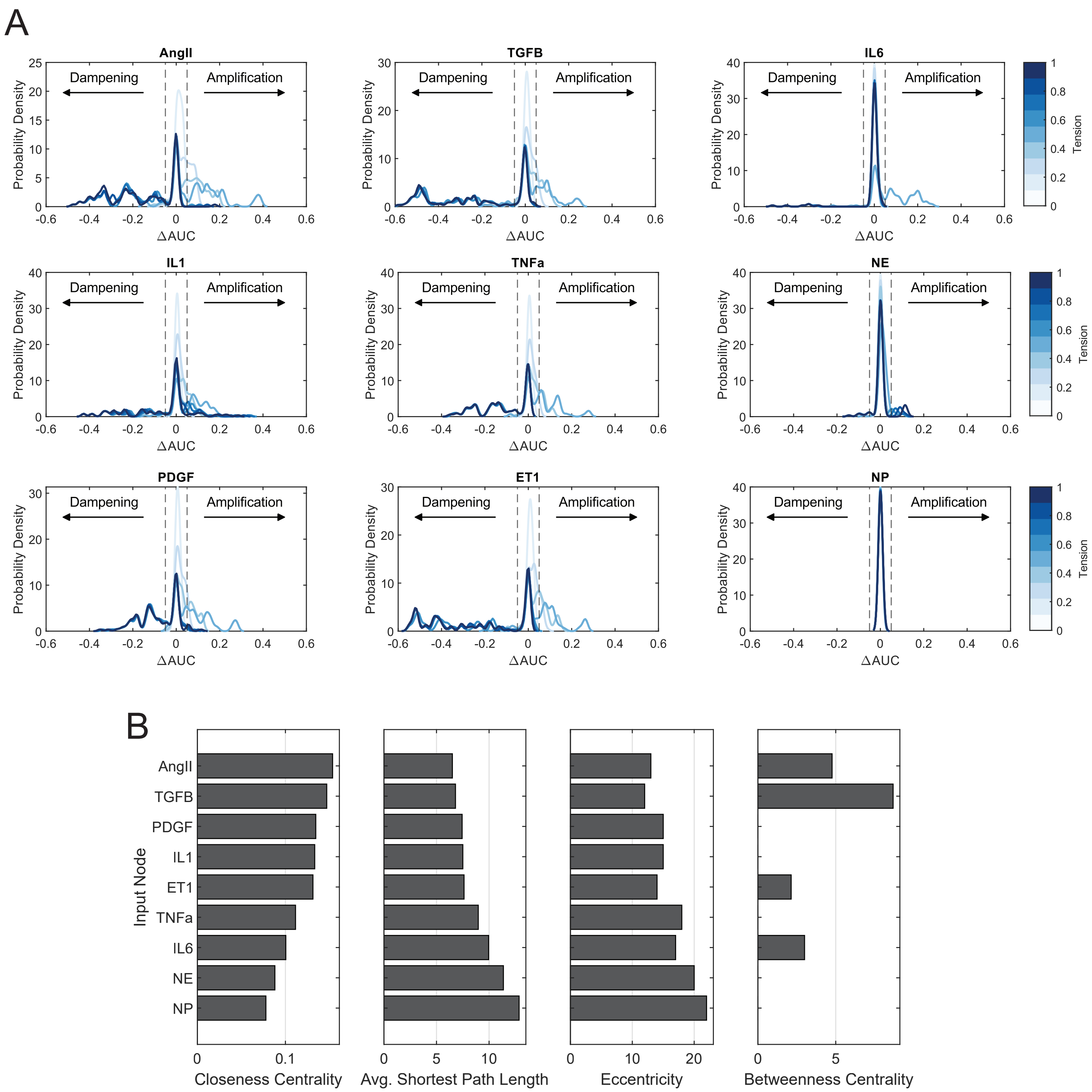


**Figure S3, Related to Figure 3. Comparison of mechano-chemo interaction trends across individual biochemical stimuli.** (A) Probability density distributions of changes in area under normalized dose-response curves (ΔAUC) with incremental levels of tension. Distributions were estimated from data using a kernel density smoothing function with a bandwidth of 0.01, and dashed lines represent a ±5% threshold used to identify nodes that exhibited amplified or dampened responses towards biochemical stimuli with added tension. (B) Comparison of topological measures of centrality for biochemical inputs. Topological analysis of the full network was conducted using the NetworkAnalyzer plugin for Cytoscape.

### Supplemental Figure 4

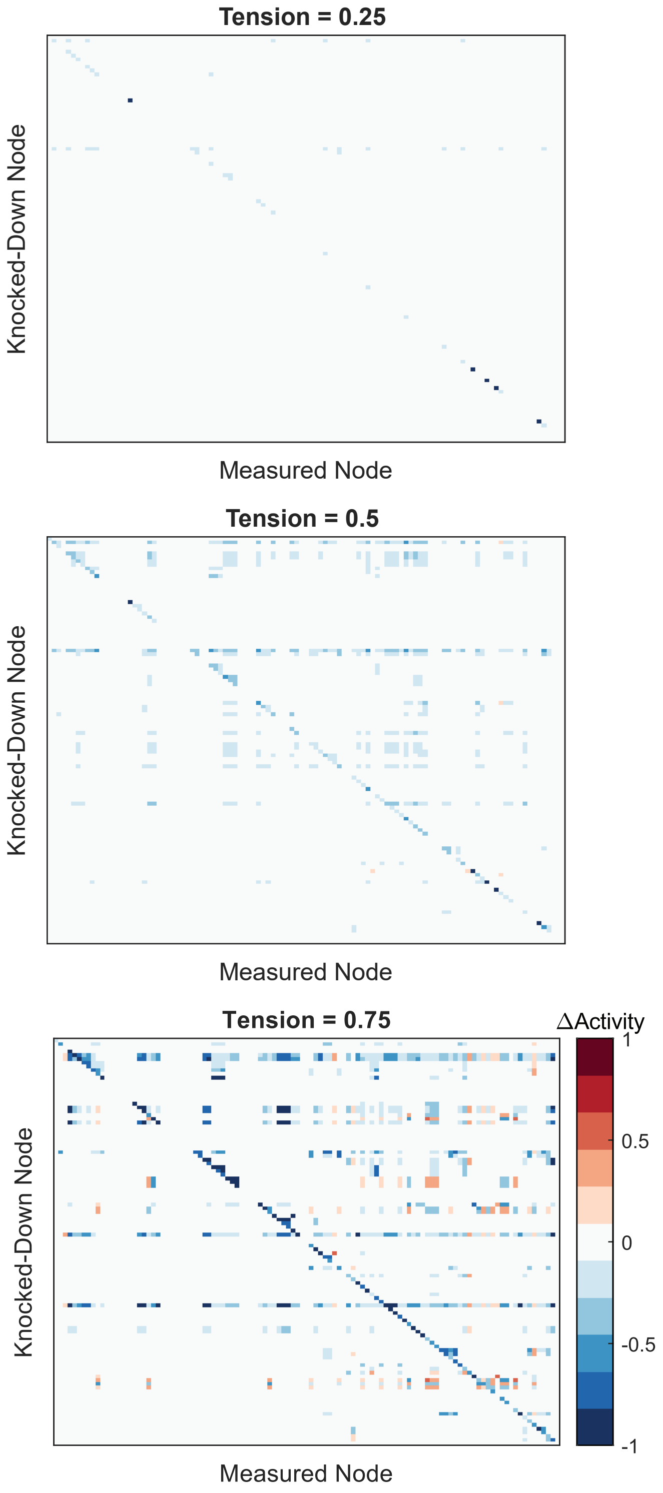


**Figure S4, Related to Figure 4. Full network perturbation anaalysis results.** Changes in activity for all nodes in the network were measured following comprehensive knockdown individual nodes ($Y_{max}$ = 0.1) under given levels of tension. Values reflect changes in node activity between perturbed and un-perturbed conditions at each tension level. Refer to Supplemental File S1 for the order of nodes perturbed/measured.

**Supplemental Figure 5**

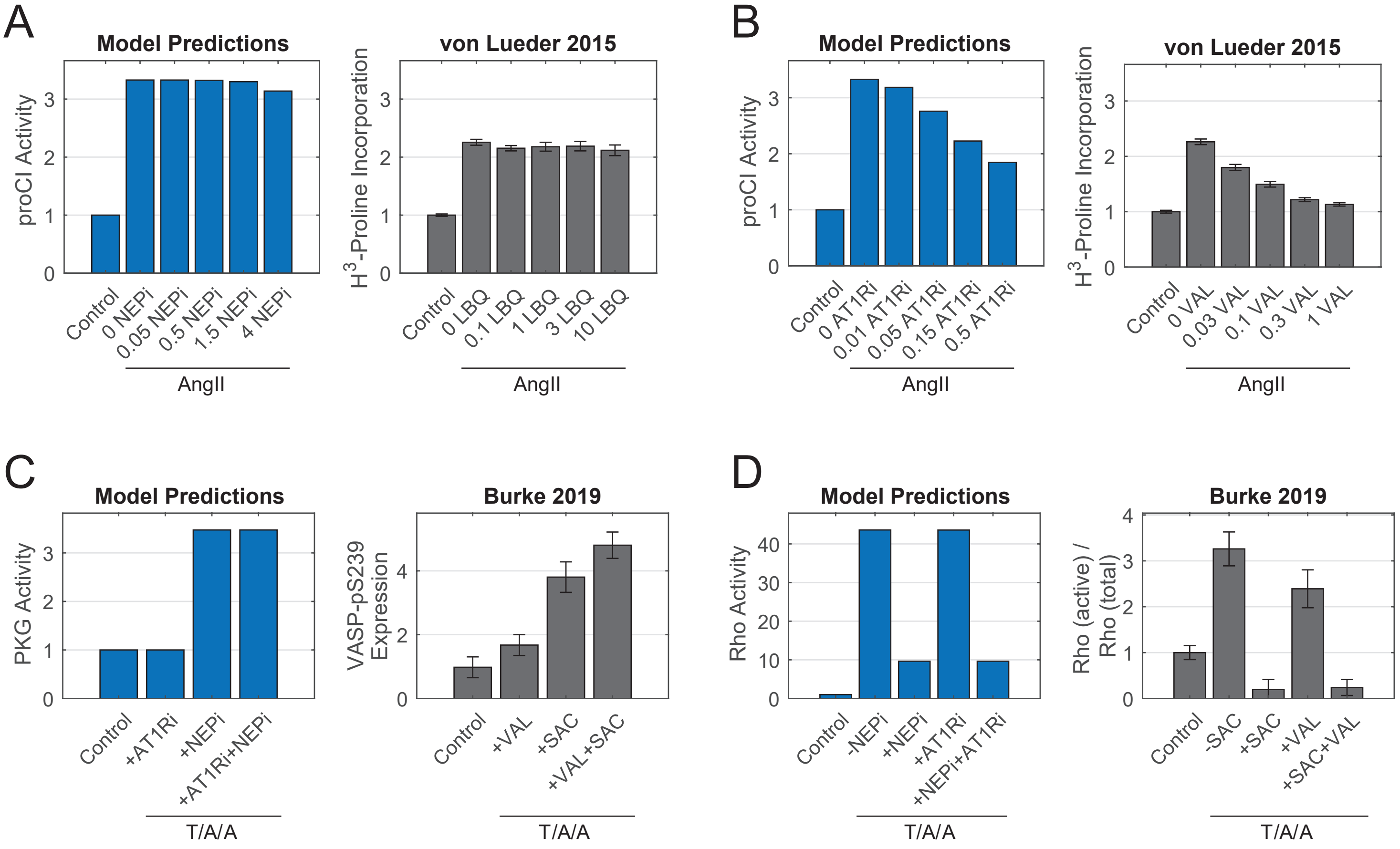


**Figure S5, Related to Figure 5. Comparisons of model-predicted and experimentally-measured changes in procollagen expression and signaling intermediate activity.** (A) Simulated and experimental measurements of procollagen I/collagen synthesis in response to increasing doses of neprilysin inhibitor LBQ657 (LBQ). Simulated concentrations of NEPi represent positive modifiers added to $Y_{max}$ parameters, and experimental concentrations represent concentrations of LBQ657 in μM. (B) Simulated and experimental measurements of procollagen I/collagen synthesis in presponse to increasing doses of angiotensin receptor blocker valsartan (VAL). Simulated concentrations represent negative modifiers subtracted from $Y_{max}$ parameters, and experimental concentrations represent concentrations of valsartan in μM. (C) Simulated and experimental measurements of PKG activity in response to valsartan and/or neprilysin inhibitor sacubitril (SAC). Experimental values reflect phorphorylation of PKG analog VASP (vasodilator-stimulated phosphoprotein). (D) Simulated and experimental measurements of Rho activity in response to valsartan and/or sacubitril. All experimental data represent mean levels ± SEM.

**Supplemental Figure 6**

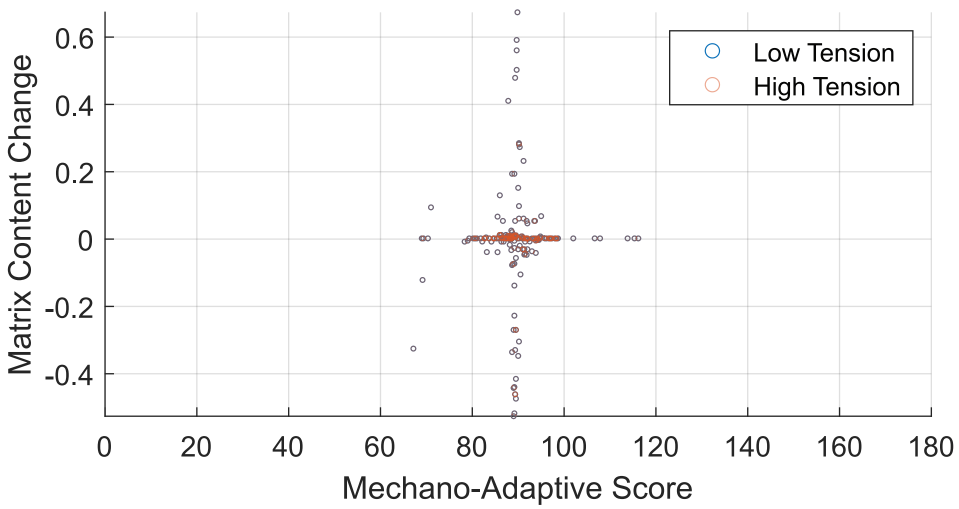


**Figure S6, Related to Figure 6. Screen of individual drug targets for mechano-adaptive matrix expression.** Changes in matrix content as a function of output node expression were measured following comprehensive knockdown and overexpression of individual nodes in low or high tension contexts. Perturbations were scored for mechano-adaptive behavior based on matrix content change values as described in STAR Methods, Mechano-Adaptive Drug Screen section. Overlapping low- and high-tension values indicate perturbations that equally altered matrix content regardless of the tensional context.

**Table S1**

| Input | Tension Levels Compared | K-S Test p-value | Tension Levels Compared | K-S Test p-value | Tension Levels Compared | K-S Test p-value |
|---|---|---|---|---|---|---|
| AngII | 0.2—0.5 | 2.2E-17 | 0.2—0.9 | 2.37E-37 | 0.5—0.9 | 1.00E-32 |
| TGFB | 0.2—0.5 | 1.88E-13 | 0.2—0.9 | 6.10E-32 | 0.5—0.9 | 5.20E-31 |
| IL6 | 0.2—0.5 | 1.87E-28 | 0.2—0.9 | 9.56E-6 | 0.5—0.9 | 9.90E-23 |
| IL1 | 0.2—0.5 | 5.38E-21 | 0.2—0.9 | 5.88E-16 | 0.5—0.9 | 5.88E-16 |
| TNFa | 0.2—0.5 | 3.17E-21 | 0.2—0.9 | 1.55E-39 | 0.5—0.9 | 1.55E-39 |
| NE | 0.2—0.5 | 1.07E-22 | 0.2—0.9 | 1.19E-3 | 0.5—0.9 | 7.47E-13 |
| PDGF | 0.2—0.5 | 5.38E-21 | 0.2—0.9 | 6.10E-32 | 0.5—0.9 | 6.10E-32 |
| ET1 | 0.2—0.5 | 1.15E-15 | 0.2—0.9 | 8.30E-34 | 0.5—0.9 | 1.70E-32 |
| NP | 0.2—0.5 | 1.18E-17 | 0.2—0.9 | 2.02E-12 | 0.5—0.9 | 6.97E-4 |

**Table S1, Related to Figure 3.** Results of Kolmogorov-Smirnov tests with Benjamini-Hochberg correction comparing ΔAUC distributions for biochemical inputs between levels of tension.
